# A long non-coding RNA controls parasite differentiation in African trypanosomes

**DOI:** 10.1101/2020.05.03.074625

**Authors:** F. Guegan, F. Bento, D. Neves, M. Sequeira, C. Notredame, L.M. Figueiredo

## Abstract

*Trypanosoma brucei* causes African sleeping sickness, a fatal human disease. Its differentiation from replicative slender form into quiescent stumpy form promotes host survival and parasite transmission. Long noncoding RNAs (lncRNAs) are known to regulate cell differentiation. To determine whether lncRNAs are involved in parasite differentiation we used RNAseq to survey the *T. brucei* lncRNA gene repertoire, identifying 1,428 previously uncharacterized lncRNA genes. We analysed *grumpy*, a lncRNA located immediately upstream of an RNA-binding protein that is a *key* differentiation regulator. Grumpy over-expression resulted in premature parasite differentiation into the quiescent stumpy form, and subsequent impairment of *in vivo* infection, decreasing parasite load in the mammalian host, and increasing host survival. Our analyses suggest Grumpy is one of many lncRNA that modulate parasite-host interactions, and lncRNA roles in cell differentiation are probably commonplace in *T. brucei*.

## Introduction

When *T. brucei*, a unicellular kinetoplastid parasite, reaches a critical density in the mammalian blood, a quorum-sensing mechanism is activated and the parasites differentiate into quiescent, non-dividing stumpy forms (Vassella et al., 1997), limiting parasite population size and extending host survival. The stumpy form also facilitates transmission to the tsetse fly vector and development into insect procyclic forms (Silvester et al., 2017). In *T. brucei*, parasite density is sensed via the stumpy induction factor (SIF) (Vassella et al., 1997) and the SIF signaling pathway, which promotes gene expression, morphological, and metabolic changes associated with the stumpy form (Mony et al., 2014). To date, 43 genes have been identified to function in the SIF signaling pathway, ranging from signal transduction to signal response (Mony et al., 2014). RBP7 RNA-binding proteins (RBP7A and RBP7B) are effectors molecules of this pathway controlling downstream gene expression. RBP7A/B null mutant parasite become unresponsive to SIF signal and are unable to differentiate into stumpy forms. RBP7 genes are therefore key regulators of parasite differentiation, yet their mode of action and target genes are unknown (McDonald et al., 2018; Mony and Matthews, 2015).

In eukaryotes, lncRNA gene abundance is comparable to that of protein coding genes (Rinn and Chang, 2012). LncRNAs function in many cellular pathways (Carlevaro-Fita and Johnson, 2019; Geisler and Coller, 2013; Yao et al., 2019), including cell differentiation (Flynn and Chang, 2014; Ransohoff et al., 2018). LncRNAs can regulate cell fate choice by either promoting or inhibiting differentiation. In skin stem cells, ANCR (anti-differentiation noncoding RNA) and TINCR (terminal differentiation noncoding RNA) lncRNAs function antagonistically. While ANCR suppresses epidermal differentiation pathway and maintains the stem cell compartment, TINCR promotes epidermal terminal differentiation (Kretz et al., 2013, 2012). LncRNAs are also important players in parasite infections, regulating antigenic variation of *Plasmodium falciparum* (Amit-Avraham et al., 2015; Guizetti et al., 2016) and associated to the host cell response in *Toxoplasma gondii* infection (Menard et al., 2018).

To date, only 95 putative lncRNA genes have been annotated in *T. brucei*, all with unknown functions (Kolev et al., 2010). This small number, compared to 9598 *T. brucei* protein coding genes (Aslett et al., 2009), prompted us to analyze the non-coding repertoire of *T. brucei* and to determine if lncRNAs are regulators of parasite differentiation.

## Results and Discussion

We used a combination of strand-specific and paired-end RNASeq, in-silico analysis, and database integration to re-annotate the lncRNA gene repertoire of *T. brucei* (Figure 1A - figure supplement 1-5). We identified 1,428 previously uncharacterized transcripts longer than 200 nt, having no significant coding potential, few ribosomal interactions, and which do not encode any unique peptides (table supplement 1 and 2 - figure supplement 2-5). These putative lncRNAs are scattered throughout the 11 chromosomes of the *T. brucei* genome in a mostly intergenic fashion (figure supplement 6). They are shorter, less expressed, and less GC-rich than *T. brucei* protein coding mRNAs (figure supplement 7) but they otherwise harbor regular mRNA trans-splicing/polyadenylation motifs (figure supplement 8). We detected these transcripts either in the nucleus and/or the cytoplasm of various *T. brucei* life cycle stages (Figure 1B). In total, 25% of the lncRNA transcripts are differentially expressed between mammalian bloodstream and insect procyclic forms (figure supplement 9 - table supplement 3), compared to 16% of differentially expressed proteins coding transcripts. LncRNA gene repertoire in *T. brucei* is substantial (11% of total genes) and shows a high dynamic expression pattern during the parasite life cycle.

We analyzed an RNAi screen data output (Alsford et al., 2011) to test our hypothesis that lncRNAs are involved in parasite differentiation. We found a total of 399 lncRNA genes that appear to be required for differentiation to occur (Figure 1C - table supplement 4), consistent with our expectation that *T. brucei* lncRNA regulate parasite transition and adaptation between mammalian and insect vector hosts. LncRNAs have been reported to regulate cell differentiation by modulating expression of their neighboring genes (Flynn and Chang, 2014). We found 19 *T. brucei* lncRNAs genes located immediately upstream or downstream of 18 of the 43 SIF pathway genes (table supplement 5). The lncRNA Ksplice-3137a, which we named *grumpy* (for regulator of Growth and Stumpy formation), is located upstream of RBP7A and RBP7B, which are both required for SIF-induced stumpy formation (Mony et al., 2014). *grumpy’s* pattern of expression is similar to that of RBP7, which is transcribed both in the bloodstream and procyclic forms of *T. brucei* (Figure 2A). However, unlike RBP7 transcripts, *grumpy* does not interact with *T. brucei* ribosomes (Figure 2A) and does not produce detectable peptides (table supplement 2).

**Figure 1.**
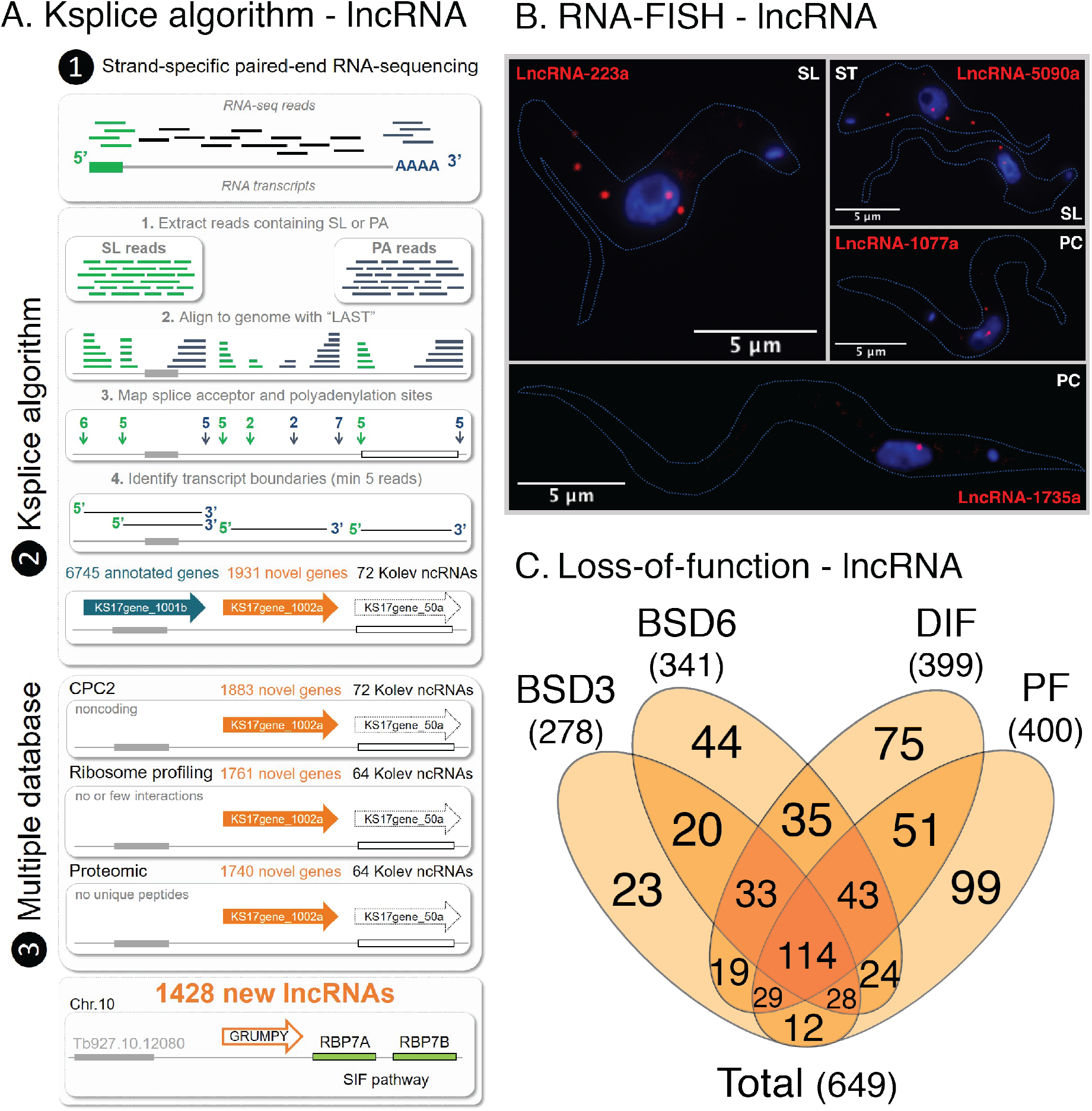
Identification of 1,428 lncRNAs in *T. brucei*. (A) Pipeline used for the identification of lncRNAs genes in *T. brucei*. (1) Strand-specific and paired-end RNA-seq. (2) Ksplice identified putative genes whose transcripts contained a spliced-leader sequence (SL) and a poly(A) tail (PA) at the extremities. Ksplice used LAST (Kielbasa et al., 2011) to map RNA-seq reads to *T. brucei* genome. (3) The non-coding nature of the putative lncRNAs was predicted from a low CPC score, poor association with ribosomes, and no detectable peptides. *Grumpy* lncRNA is intergenic and immediately upstream of RBP7 genes, previously shown to be involved in SIF-dependent pathway. (B) Subcellular localization of Ksplice long noncoding RNA genes in slender, stumpy, and procyclic forms of *T. brucei*, using RNA-FISH. (C) Number of Ksplice lncRNA genes that causes loss of parasite fitness upon downregulation by RNA interference (extracted from RIT-seq analysis (Alsford et al., 2011)). RNA interference was induced in bloodstream forms for 3 days (BSD3) — 278 lncRNAs; in bloodstream forms for 6 days (BSD6) — 341 lncRNAs; during *in vitro* parasite differentiation from bloodstream to insect procyclic forms (DIF) — 400 lncRNAs; in procyclic forms, PF — 402 lncRNAs. The total number of lncRNA genes essential for parasite fitness in this screen was 649.

**Figure 2.**
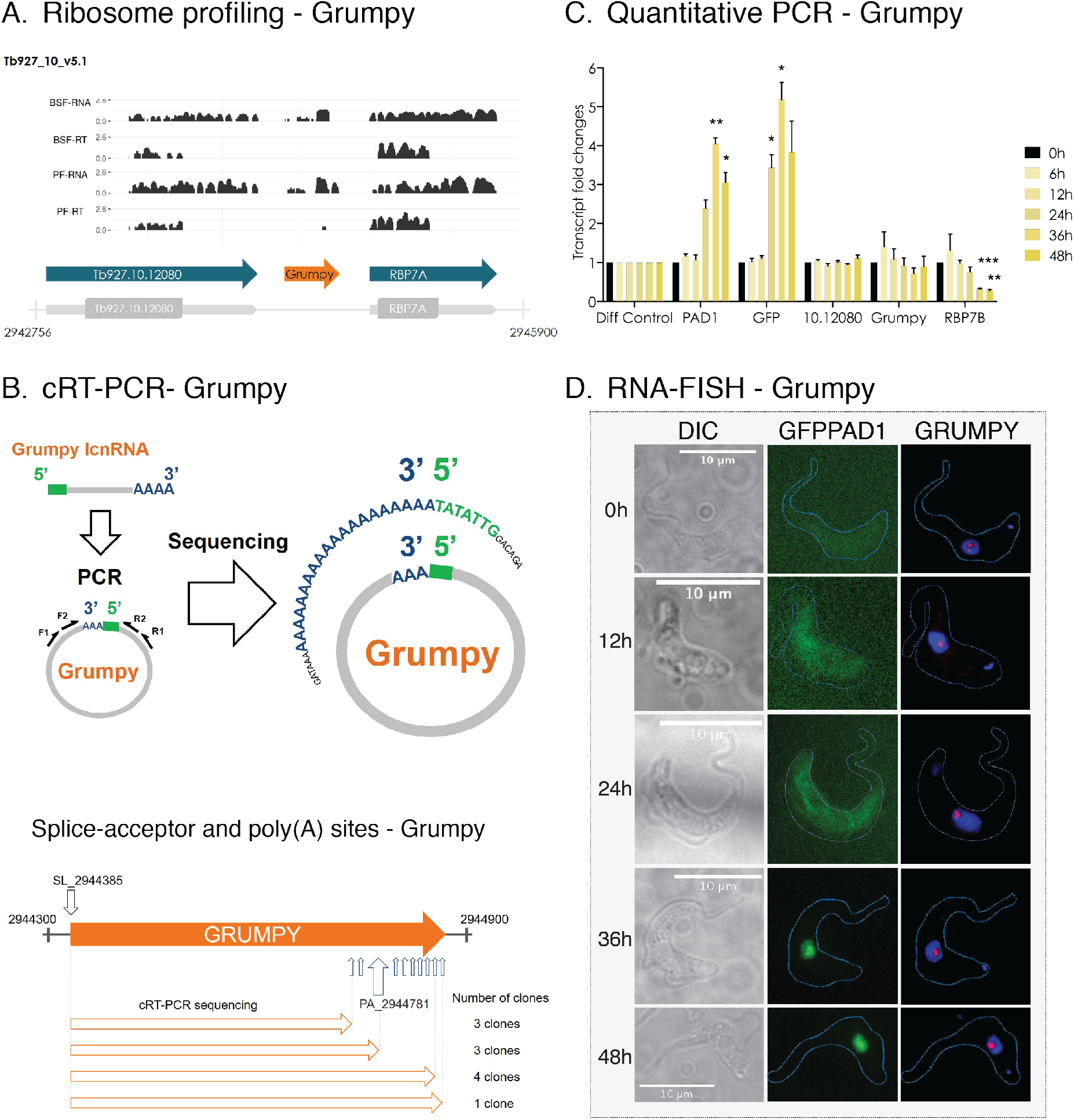
Dynamic subnuclear localization of *grumpy* during parasite differentiation. (A) Ribosome association of *grumpy* and its neighboring genes was assessed by analyzing previously published ribosome profiling datasets: Mapping of RNA-seq reads from bloodstream forms (BSF-RNA) or procyclic forms (PF-RNA); mapping of ribosome profiling reads from bloodstream forms (BSF-RT) or procyclic forms (PF-RT). (B) Sequencing and accurate mapping of the 5’ and 3’ ends of the *grumpy* lncRNA using circular RT-PCR (cRT-PCR). Black outlined arrows show the position of splice acceptor site (SL) and polyadenylation sites (PA) identified with our Ksplice algorithm in the *grumpy* gene locus. Orange outlined arrows show the *grumpy* transcript isoforms that we sequenced using cRT-PCR and the number of clones sequenced for each isoform. (C) Transcript level changes during the transition from slender to stumpy forms, measured by qRT-PCR. Stumpy formation was induced by pCPT-cAMP (C3912 Sigma-Aldrich). Diff (Tb927.10.12970) is used as control to normalize transcript levels (Barquilla et al., 2008). PAD1 and GFP genes are used as controls to estimate parasite differentiation into stumpy forms. 10.10280 (Tb927.10.12080) is the gene upstream of *grumpy*. Results are shown as mean (SEM, n=3). Dunnett’s multiple comparisons test is used for statistical analysis using the time point 0h as the control for comparison (Adjusted P value: * <0,05; ** <0,01; *** <0,001). (D) Subcellular localization of *grumpy* during the transition from slender to stumpy forms using RNA-FISH. Time points (0h, 12h, 24h, 36h, 48) of parasite differentiation after addition of the pCPT-cAMP stimulus to the culture medium. DIC: Differential interference contrast microscopy image of *T. brucei;* GFP: GFP::PAD1 signal expressed in nucleus of stumpy forms; *GRUMPY: grumpy* signal using RNA-FISH (Stellaris probes).

To further characterize the *grumpy* transcript, we used a circular RT-PCR (cRT-PCR) assay, in which *T. brucei* RNAs are circularized via their 5’ – 3’ end junctions, amplified and sequenced. We used gene-specific primers to confirm that *grumpy* is a trans-spliced and polyadenylated lncRNA transcript expressed as at least five different isoforms, including the smallest (359bp), the major (397 bp) and longest forms (432 bp) (Figure 2B). These findings are consistent with the Ksplice in-silico analysis, which revealed one splice-acceptor site and 10 alternative polyadenylation sites for *grumpy* (Figure 2B). RT-qPCR showed that, while the mRNA levels of RB7B decrease during parasite differentiation from slender to stumpy forms, *grumpy* levels remain constant (Figure 2C). However, an RNA-FISH analysis revealed changes in the subcellular localization of *grumpy* during stumpy formation (Figure 2D). Whereas in slender forms *grumpy* localizes in three distinct nuclear foci (one in the nucleolus and two in the nucleoplasm), in stumpy forms *grumpy* localizes in a single nucleolar focus (Figure 2D). Contrary to our initial hypothesis, these changes in subcellular localization suggest that *grumpy* may act through a trans-acting mechanism.

To identify the function of *grumpy*, we used a gain-of-function approach, in which *grumpy* was over-expressed 3-fold from an exogenous genomic location (mini-chromosome). These exogenous *grumpy* transcripts retained the original nucleolar localization and the transcript levels of RBP7A and B remained unchanged (Figure 3A and B). We observed that exogenous expression of *grumpy* repressed *T. brucei* growth and increased lifespan *in vitro* (Figure 3C and D). We asked whether this reduction in parasite growth could be explained by a higher proportion of the stumpy forms in culture. Stumpy formation occurs only at high parasite density via the SIF-dependent quorum-sensing mechanism, and can be quantified using flow cytometry (Batram et al., 2014; Dean et al., 2009) by measuring the fraction of transgenic parasite expressing the fluorescent stumpy marker GFP::PAD1. After 2 days in culture, 60% of the *grumpy* over-expressing parasites with were in the stumpy form, compared to 7% in the parental line cultured for the same time period (Figure 3E). *Grumpy* over-expression also lead to a lower parasite density (<0.7×10^6^ cells/ml) compared to the parental culture (1.4×10^6^ cells/ml) (Figure 3C). Parasites over-expressing *grumpy* displayed all hallmarks of being in stumpy form, including PAD1 protein expression at the cell surface (Figure 3F), arrest at the cell cycle G0/G1 phase (Figure 3G), and pre-adaptation to differentiate into the insect procyclic stage (Figure 3H).

**Figure 3.**
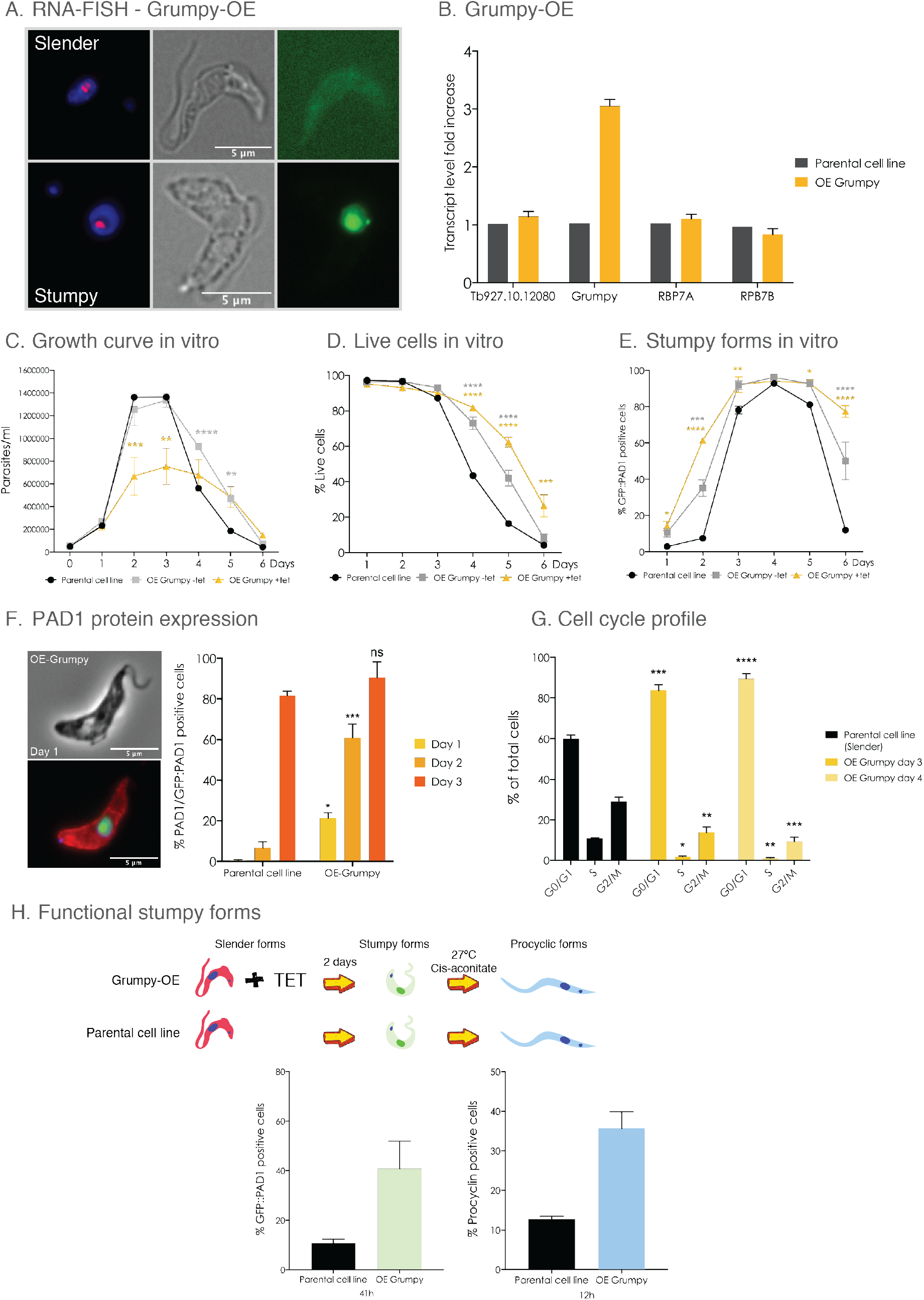
*Grumpy* over-expression promotes premature parasite differentiation. (A) Subcellular localization of *grumpy* after over-expression using RNA-FISH. Left-to-right: *GRUMPY: grumpy* signal using RNA-FISH; Phase contrast signal of *T. brucei* parasite; GFP: GFP::PAD1 signal expressed in the nucleus of stumpy forms. (B) Transcript levels measured by qRT-PCR of *grumpy* and its neighbouring genes in parental cell line (black bars) and in *grumpy* over-expressing cell line (yellow bars). Changes of transcript levels were measured by normalizing transcript level to a control gene (Tb927.10.12970) and to the parental cell line. (C) Growth in parasites over-expressing *grumpy* (without passage) for 6 days. (D) Percentage of live cells measured by FACS of propidium iodide-stained cells after inducing *grumpy* overexpression. (E) Percentage of GFP::PAD1 positive parasites (stumpy forms) measured by FACS after inducing *grumpy* over-expression. (F) Percentage of parasites expressing both GFP::PAD1 and endogenous PAD1 protein are measured by microscopy and image quantification, after inducing *grumpy* over-expression. Microscopy picture (on the left) showed an example of parasite expressing both GFP::PAD1 in the nucleus (in green) and the endogenous PAD1 protein at the cell surface (in red). Parasite DNA stained with DAPI (in blue). (G) Cell cycle profile of parental cell line (slender forms) and *grumpy*-over-expressing parasites at day 3 and 4 of *in vitro* culture without passage. (H) Differentiation assay to separately follow the transition from slender to stumpy and stumpy to procyclic forms. Parasites were cultured for 2 days without passage, in the presence or absence of tetracyclin. GFP::PAD1 expression was measured by FACS to score the percentage of stumpy forms. Parasites were transferred to DTM medium containing cis-acconitate and placed for 12h at 27°C. The percentage of procyclic forms was quantified by flow cytometry using procyclin antibody. Results from Panel B to H are shown as mean (SEM, n=3). Sidak’s multiple comparisons test is used for statistical analysis using the parental cell line as the control for comparison (Adjusted P value: * <0,05; ** <0,01; *** <0,001, **** <0,0001).

To confirm these results *in-vivo*, we induced mouse infections with parasites overexpressing *grumpy* and measured parasitemia, mouse survival, and stumpy formation, which were compared to infection by the parasite parental line. Mice infected with the parental cell line showed a typical infection profile characterized by successive waves of parasitemia (Figure 4A) and an average survival of 43 days (Figure 4B) (Trindade et al., 2016). By contrast, mice infected with parasites over-expressing *grumpy* showed no detectable parasitemia and did not die from the infection (>100days) (Figure 4A and B). When *grumpy* over-expression was induced four days post-infection, the parasites succeeded in establishing an infection (Figure 4A), with three mice out of four dying from the infection and mice survival time increasing from approximately 43 to 72 days (Figure 4B). Thus, *grumpy* over-expression substantially reduces parasite virulence in mice.

**Figure 4.**
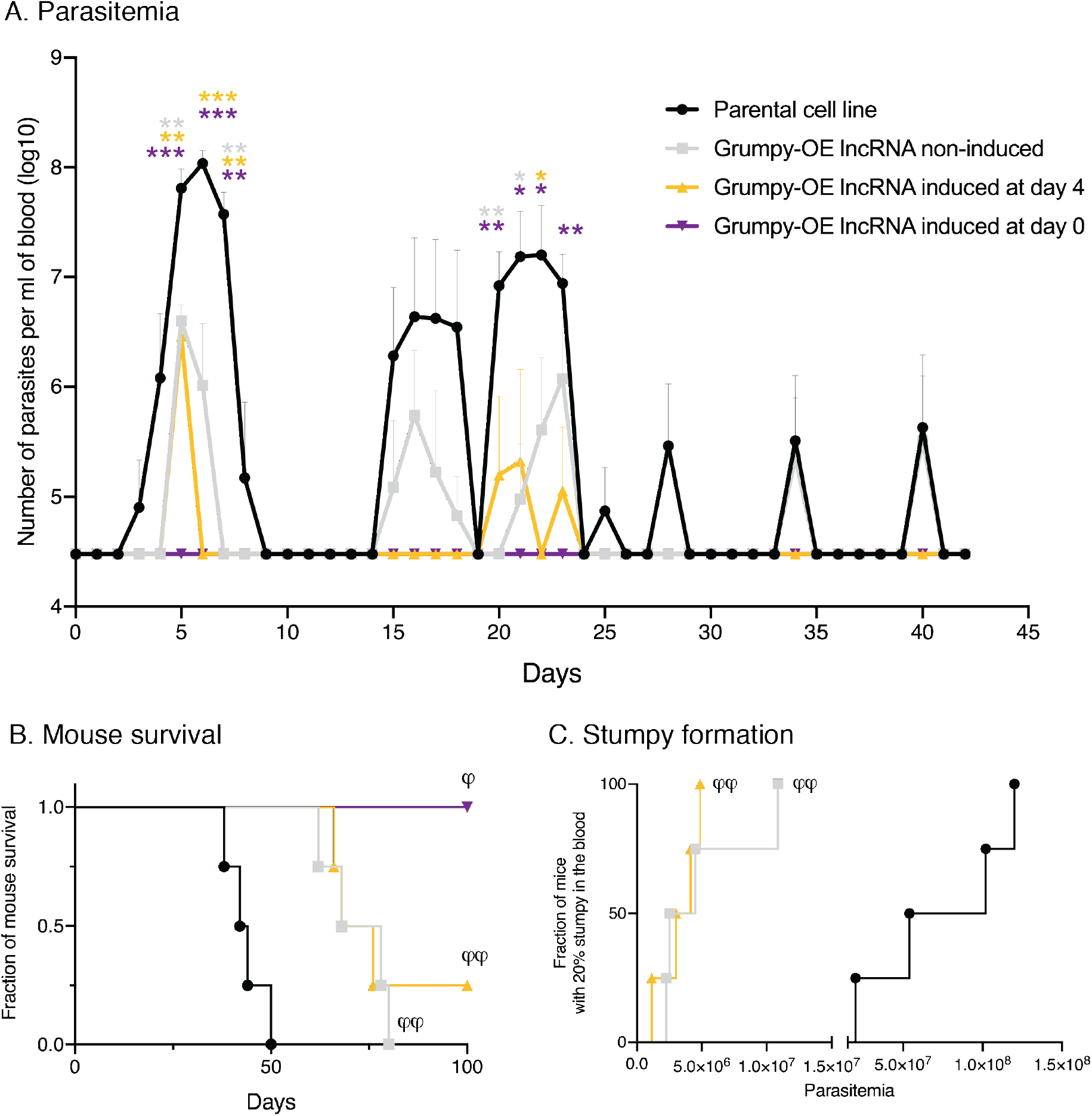
Over-expression of *grumpy* promotes premature differentiation into stumpy forms *in vivo* and prolongs mouse survival. (A) Parasitemia in mice infected with the parental cell line (black line) or with a *grumpy*-over-expressing cell line. *Grumpy* over-expression was either not induced or induced by adding doxycycline to drinking water either at Day 0 (purple curve) or Day 4 (yellow curve) of infection. Results are shown as the mean (SEM, n=4). Dunnett’s multiple comparisons test is used for statistical analysis using the parental cell line as the control (Adjusted P value: ** <0,01; *** <0,001). (B) Mice survival rates according to the type of infection described in panel A. Log rank (Mantel-Cox) test for comparisons of Kaplan-Meier survival curves indicates a significant increase in the survival rates in mice infected with *grumpy*-over-expressing cell-line parasites compared to mice infected with the parental cell line. φ φ, *p*=0.0067. φ, *p*=0.0177. (C) Fraction of mice with at least 20% of GFP::PAD1 positive parasites (=stumpy forms) in the blood as a function of the parasitemia. Log rank (Mantel-Cox) test for comparisons of Kaplan-Meier curves indicates a significant premature parasite differentiation into stumpy forms in mice infected with *grumpy*-overexpressing cell line parasites compared to mice infected with the parental cell line. φ φ, *p*=0.0067.φ, *p*=0.0177.

Our *in vivo* analysis also recapitulated *in vitro* observations with respect to stumpy forms and density. Wild-type parasites started differentiating into stumpy forms (>20% stumpy forms in the blood) only at high parasitemia (>1.5×10^7^ parasites/ml) whereas *grumpy*-over-expressing parasites differentiated into stumpy forms when parasitemia was as low as 1.1×10^6^ parasites/ml (Figure 4C). These results support the notion that *grumpy* over-expression triggers premature *T. brucei* differentiation into stumpy forms, which is associated with a reduction in parasite virulence.

Here, we show that the *grumpy* lncRNA is a regulator of parasite differentiation in *T. brucei*. Its mechanism of action, currently unknown, does not correlate to RBP7 expression but likely involves its nucleolar localization. This localization is seen for the mammal X inactivation factor, Xist, and pRNA, a promoter-associated RNA that drives mouse embryonic stem cell differentiation (Savić et al., 2014; Zhang et al., 2007). Both of these lncRNAs promote heterochromatin formation at the nucleolar periphery. Other lncRNAs bind and sequestrate specific proteins in the nucleolus, rendering them functionally inert (Audas et al., 2012). *Grumpy* could regulate stumpy formation through similar mechanisms either by sequestering stumpy regulator proteins (or its mRNAs) or by modulating the genomic conformation of stumpy regulator genes within the nucleolus. In the future, it will also be important to evaluate if grumpy is necessary for stumpy formation. This will require careful genome editing given that part of the grumpy lncRNA shares sequence homology with RBP7A 3’UTR.

Of the 1,428 lncRNA genes we have identified in *T. brucei*, 649 have been predicted, via an RNAi screen, to play a role in parasite fitness, including 399 lncRNA genes involved in cell differentiation (Figure 1C). That 18 of the 41 genes involved in the quorum-sensing signaling pathway have a lncRNA gene in close proximity suggests that *grumpy’s* role in differentiation may be only one instance of a more general process used by the parasite to sense its environment and modulate its virulence accordingly. Understanding these regulatory processes may open up possibilities for developing therapeutic strategies to treat sleepiness sickness.

## Materials and Methods

### Ethics statement

Male C57BL/6J (6–8 weeks old) were purchased from Charles River Laboratories (Lyon, France). All animal care and experimental procedures were performed according to EU regulations (Directive 2010/63/EU9) and approved by the Animal Ethics Committee of Instituto de Medicina Molecular João Lobo Antunes (AWB2016_19FG_RNA).

### *T. brucei* cell culture

A stumpy reporter cell line with a GFP:PAD1UTR construct (Batram et al., 2014) integrated into the tubulin locus was generated in *T. brucei* Antat1.1e (90:13) strain (Engstler and Boshart, 2004). The stumpy reporter cell line was selected for its most intense GFP expression, which occurs in the nucleus in response to quorum-sensing signal. This reporter cell line was used as the genetic background to overexpress for Grumpy-lncRNA. It was cultivated in HMI-11 at 37°C in 5% CO_2_ with 2,5 μg/mL G418, 5 μg/mL Hygromycin B, 5 μg/mL Blasticidin S and 2,5 μg/mL Phleomycin.

### Grumpy-lncRNA expression construct

The non-coding sequence of Grumpy lncRNA was amplified from *T. brucei* Antat 1.1E genomic DNA with forward (5’-CAAAAGGACAGAATTATAGGTTCA-3’) and reverse (5’-GATGCAGCTCAACAGCAAG-3’) primers and inserted into pDEX577 (phleo) between the Hind III and BamH I sites of the plasmid. pDEX577 vectors are highly-modular expression vectors for inducible expression of transgenes, integrating in the minichromosome repeats, which was designed and constructed by Steve Kelly (Kelly et al., 2007). Moreover, two T7 terminator sequences were inserted between the BamH I and Kpn I sites of the plasmid just downstream to the Grumpy lncRNA construct. The construct was linearized with Not I prior to transfection. Stable transfectant clones were obtained by serial dilution of the transfected population and selected after 6-7 days after transfections. Inducible expression is obtained by adding Tetracycline (in vitro) or Doxycycline (in vivo) at the following concentrations: 1 μg/mL and 1 mg/ml.

### Inducible expression of Grumpy-lncRNA

Cells were diluted at 5×10^4^ parasites/mL and induced with 1 μg/mL of tetracycline for 6 days. Cells were counted every day, live/dead cells were assessed by Propidium iodide staining, GFP::PAD1 positive cells were scored and all these parameters quantified using Accuri C6 flow cytometry. At day 2 after tetracycline induction, RNA samples were collected by centrifugation of equivalent number of cells and addition of TRIzol reagent (Invitrogen) to the cell pellets. At day 3 after tetracycline induction, an equivalent number of cells were collected by centrifugation and the cell-cycle profiles were assessed using Accuri C6 flow cytometer and Propidium iodide staining in fixed cells.

### Quantitative RT-PCR

RNA was prepared with TRIzol reagent (Invitrogen) according to manufacturer’s instructions and cDNA was synthesized with random primers and SuperScript II reverse transcriptase. Quantitative PCR was performed with AmpliTaq Gold™ DNA Polymerase (Power SYBR Green Master Mix, Applied Biosystems™) and gene-specific primers:

Control of Differentiation (Tb927.10.12970)

FW: CCAGCCTTCTCAATCTCCAG

Rv: GGCCACAGTTGGATAGCTTG

Tb927.10.12080

FW: CCTGCAGGCGTCACATTC

RV: CAGTGAAGAAGAAAAGGCACG

Grumpy lncRNA:

FW: AACGGAAGGAAAGTTTGTGAATGC

Rv: GTGAATGAACTTTTTGTTTGGCGTC

RBP7A:

FW: GCTCGACTTTTTGTTGGGCAG

RV: CATATTGTAGCGGTTGTGAAGCG

RBP7B:

FW: CTTTAACGCAACCGAAGATG

RV: CAACGGTTGTGAAGTCCG

The quantitative PCR program was:

Stage 1 - 10 min at 95°C

Stage 2 - 15 sec at 95°C, 15 sec at 60°C, 30 sec at 72°C (40 cycles)

Melt curve - 15 sec at 95°C, 1 min 10 sec at 60°C, 15 sec at 95°C

### Stumpy formation assay

Cell cultures were started at 5×10^4^ parasite/mL and induced or not with 1 μg/mL of tetracycline. Every day of culture, sufficient number of cells (>10 000 parasites) were collected, washed with trypanosome dilution buffer (TDB) (5 mM KCl, 80 mM NaCl, 1 mM MgSO4, 20 mM Na2HPO4, 2 mM NaH2PO4, 20 mM glucose, pH 7.4), and resuspended in 200 μL of TDB with 1 μg/mL Propidium iodide. A fixed volume of each cell culture was analysed by flow cytometry (Accuri C6) to simultaneously measure the parasites density, live and dead parasites and the GFP::PAD1 expression.

### Cell cycle profile assay

Cell cultures were started at 5×10^4^ parasite/mL and induced or not with 1 μg/mL of tetracycline. After 3 and 4 days of *in vitro* culture, 2×10^6^ parasites were collected and span down (10 min, 1300g, 4°C), washed once with ice-cold PBS, resuspended in 1 ml PBS/2 mM EDTA and fixed by adding drop wise 2.5 ml ice cold 100% ethanol (store EtOH at −20°C). Cells were fixed at 4°C for at least one hour, washed once with 1 ml PBS/EDTA at RT and resuspended in 1 ml PBS/EDTA. RNA was digested by adding 1 μl RNaseA (10 μg/μl) and DNA stained by adding 1 μl propidiumiodide (1mg/μl) during 30 min at 37°C. Cell-cycle profile were analysed by flow cytometry using Accuri C6 machine with FL3 channel.

### Parasite differentiation into procyclic assay

Cell cultures of bloodstream forms were started at 5×10^4^ parasite/mL and Grumpy lncRNA was induced or not with 1 μg/mL of tetracycline. After 2 days of *in vitro* culture, the number of stumpy forms were assessed by measuring the GFP::PAD1 expression using flow cytometry (Accuri C6). Bloodstream forms culture were collected, span down, resuspended in Differentiation Trypanosome Medium (DTM) with 6mM cis-Aconitate at 1×10^6^ parasites/mL and incubated at 27°C. Parasite differentiation into procyclic forms was assessed at 12h post differentiation by Flow cytometry using anti-*Trypanosoma brucei* procyclin antibody (Clone TBRP1/247, CLP001AP, 0.5mg, Cedarlane) conjugated with Alexa Fluor 647 (Protein labelling kit, Molecular probes) (1/500 dilution in TDB).

### Infections and Sample Collection

Four weeks old male c57BL/6 mice (Charles River, France) were inoculated intraperitoneally with 2000 parasites. Mice were infected with either Antat1.1 90:13 GFP::PAD1 cell line or Antat1.1 90:13 GFP::PAD1 Grumpy-overexpression cell line. Mice infected with Antat1.1 90:13 GFP::PAD1 Grumpy-overexpression parasites were separated in three different cages, one cage of 4 mice received only water, one cage of 4 mice received water with 1 mg/ml doxycycline hyclate (Sigma-Aldrich) at day 4 post infection, one cage of 3 mice received water with 1 mg/ml doxycycline hyclate at the day of infection. Parasitaemia was monitored by tail-vein bleeds every other day and counted using a Hemocytometer with 1:150 blood dilution in TDB. The percentage of stumpy forms in the mice blood were assessed by measuring GFP::PAD 1 expression in blood diluted sample using Accuri C6 flow cytometer. Mice survival were monitored every other day until 100 days post infections. Mice were euthanized at the first signs of severe disease distress, with all efforts to minimize animal suffering.

### RNA-FISH

Between 2,5×10^5^ to 1×10^6^ cells were harvested by centrifugation (10 min, 1800 g), washed with 1X PBS or TDB and resuspended in between 500 μL and 1 mL of fixation buffer (3,7% Formaldehyde diluted in RNAse-free PBS) for 10 min at room temperature. Fixed cells were washed with between 500 μL and 1 mL of RNAse-free PBS and resuspended with 150 μL of RNAse-free PBS. Cells were then settled on pre-coated polylysine culture dishes (35mm glass bottom, MatTEK) for at least 20 min. PBS was removed and cells were permeabilized with 1 mL of ethanol 70% (in RNAse free water) for at least 1 hour at +2 to +8 °C. Ethanol 70% is discarded and cells washed with 200 μL wash buffer A (10% vol./vol. formamide in 1X Wash Buffer A, Biosearch Technologies Cat# SMF-WA1-60). Cells were incubated with 100 μL Hybridization buffer containing 1,25 μM of RNA-FISH probes in the dark at 37 °C overnight (~16 hours). Cells were washed with 200 μL of wash buffer A and incubated 200 μL of wash buffer A in the dark at 37 °C for 30 minutes. Cells were stained with a solution of 1 μg/mL of DAPI (in wash buffer A) in the dark at 37 °C for 30 minutes. Cells were washed with 200 μL of wash buffer B (Biosearch Technologies Cat# SMF-WB1-20) and incubated with it at room temperature for 2-5 minutes. 100 μL Vectashield was added to the dishes prior analysis with the Zeiss cell observer wild Field microscope.

RNA-FISH probes were designed using the online tools provide by LGC Biosearch Technologies (Stellaris Probe Designer, https://www.biosearchtech.com/support/tools/design-software/stellaris-probe-designer). 17 probes were designed for Grumpy lncRNA, and 30-43 probes for Ksplice lncRNA223a, lncRNA1077a, lncRNA1735a and lncRNA5090a.

### PAD1 staining

5×10^5^ bloodstream form parasites were harvested by centrifugation (10 min, 1800 g), washed with 1X PBS and resuspended in 500 μL of fixation buffer (4% paraformaldehyde diluted in 1X PBS) for 10 min at room temperature. Fixed cells were washed with 500 μL 1X PBS and resuspended with 100 μL of 1X PBS. Cells were then settled on pre-coated polylysine culture dishes (35mm glass bottom, MatTEK) for at least 20 min. PBS was removed and cells were permeabilized with 100 μL of 0,1% Triton in PBS for 2 min at room temperature. Permeabilized cells are washed 5 times with 200 μL of PBS and blocked with 2% BSA in PBS for 45 min at 37°C in a humidity chamber. Cells were incubated with 100 μL of the primary antibody anti-PAD1 (1/1000 in 2% BSA in PBS, antibody provided by Keith Matthews) overnight at 4°C in a humidity chamber. Cells are washed 5 times with 200 μL of PBS and incubated with 100 μL of the secondary antibody anti-rabbit (1/1000 in 2% BSA in PBS, Goat anti-Rabbit Alexa Fluor 647 #A21245 – Invitrogen) for 45 min at 4°C in a humidity chamber. Parasite DNA was stained using 100 μL DAPI or Hoechst solution (1 μg/ml) for 20 min at room temperature. Cells were washed 5 times with 200 μL of PBS and 100 μL Vectashield was added to the dishes prior analysis with the Zeiss cell observer wild Field microscope.

### Transcript quantification and Circular RT-PCR

Transcript quantification was performed by quantitative RT-PCR, as described in Aresta-Branco *et al*. (Aresta-Branco et al., 2015) except that random hexamer primers were used to generate cDNA.

Circular RT-PCR protocol was performed essentially as described in Laboratory Methods in Enzymology book (ABELSON and SIMON, 2009). Briefly, parasites were harvested by centrifugation 677 g for 10 min at 4°C and immediately resuspended in TRIzol (life technologies). Total RNA is isolated following the manufacturer’s instructions and RNA was quantified in a NanoDrop 2000 (Thermo Fisher Scientific). The ideal RNA concentration to perform the circular RT-PCR protocol is 0.5–1 mg/μl. RNA cap and poly A tail were removed by oligonucleotide-directed RNase H cleavage using Spliced-leader and oligo dT primers. After RNAse H treatment, RNA was extracted with phenol/chloroform approach and precipitated using ethanol precipitation protocol. 3-5 μg of RNAse H treated RNA was circularized using T4 RNA ligase 1 (ssRNA Ligase, New England Biolabs), RNA was extracted with phenol/chloroform approach and ethanol precipitated. RNA was reverse transcribed using gene-specific primer R1 (100 nucleotides from the 5’end of the transcript or the RNase H cleavage site) and reverse transcriptase, RT buffer and 5mM Magnesium from Superscript II kit (life technologies). The resulting cDNA molecules contain the juxtaposed 5’ and 3’ends of circular RNA. PCR is performed on the produced cDNA using gene-specific primers R2 and forward F1. R2 primer is in “nested” position compared to R1 primer and contributes to the specificity of PCR amplicon. PCR#1 product was purified using Minielute PCR purification kit (Qiagen) and a second round of PCR amplification was performed with gene-specific primers R2 and forward F2. F2 primer is in nested position compared to F1 primer position and contributes to PCR amplicon specificity. PCR#2 product was ligated to pGEM-T easy vector or TOPO vector following the manufacturer’s instructions (Promega). After transformation in bacteria and plasmid amplification, the subcloned PCR#2 fragments were amplified and sequenced using T7 and SP6 primers.

### RNA-Sequencing

*T. brucei* bloodstream-form (BSF) and procyclic-form (PF) parasites (strain Lister 427, antigenic type MiTat 1.2, clone 221a), from PL1S cell line (Yang et al., 2009), were used to generate strand-specific libraries following the manufacture instructions (Encore^®^ Complete RNA-Seq Library Systems, NuGen) for Illumina next-generation paired-end sequencing. RNA-sequencing were performed Genomics Core Facility, EMBL Heidelberg. The sequence data from this study have been submitted to the NCBI Sequence Read Archive – SRA………

### Reconstruction of *T. brucei* transcriptome

In *T. brucei*, all mature mRNAs are trans-spliced and polyadenylated which means that all mRNA transcripts start with a conserved spliced-leader sequence and finish with poly(A) tail sequence^31^. We hypothesized that any new *T. brucei* transcripts including noncoding RNA transcripts will bear these features. RNA-seq reads were assessed for quality using FastQC. In order to improve genome mappability, RNA-seq reads size were increased, if possible, by merging the paired-end reads using PEAR software - Paired-End reAd mergeR (https://cme.hits.org/exelixis/web/software/pear/). Merged and forward unmerged reads containing a minimum of 8 bp matching the SL sequence on their 5’ end were extracted for 5’ splice-acceptor site detection and the SL sequenced removed from the read. Reads containing stretches of at least 9 A’s in the merged reads, or 9 T’s in the unmerged reverse reads were extracted for poly-A site identification and and the poly-A tails removed from the read.

SL and polyA reads were aligned to *T. brucei* genome (https://tritrypdb.org/tritrypdb/; genome annotation: version v5.1) using LAST (version 959) alignment tools (Kielbasa et al., 2011) (http://last.cbrc.jp/). 5’ splice-acceptor sites were determined by the first position of all SL-containing reads mapping uniquely to the genome. Poly-A sites were determined by the last position of all uniquely mapped poly-a containing reads. SL acceptor or polyA sites were considered for further analysis if a splice-acceptor or polyA site is supported by at least 5 reads. Putative *T. brucei* genes were defined by all genomic regions separated by at least one 5’ acceptor site and one 3’ poly-A site occurring before the next downstream 5’ site. For each gene region, the longest transcript isoform was defined by the association of the most upstream SL-acceptor site and the most downstream polyA site. In contrary, the major isoform of *T. brucei* gene transcript was defined by the gene region bordered by the major SL acceptor and polyA sites (*i.e*. ones with most reads aligned). This analysis identified 8,831 genes in *T. brucei* genome

### Identification of Ksplice putative new noncoding genes

A stringent selection pipeline was developed to systematically identify *T. brucei* non-coding RNAs. This pipeline aims to discard housekeeping (tRNAs, snRNAs, snoRNAs) *T. brucei* non-coding RNAs and transcripts with protein-coding potential. First, only transcripts that do not overlapped annotated protein-coding and non-coding RNA genes from Tritryp data (https://tritrypdb.org/tritrypdb/; genome annotation: version v5.1) were retained. Second, *T. brucei* transcripts with protein-coding potential were excluded. Protein-coding potential was determined by using three different approaches. 1) The protein-coding potential for each transcript was calculated using coding potential calculator score (CPC2) (Kang et al., 2017). 2) The association with *T. brucei* ribosomes and translation efficacy of each transcript was measured using the published ribosome profiling data from *T. brucei* (Vasquez et al., 2014) and re-analysing it with our Ksplice gene annotation. 3) The non-coding potential of each transcript were confirmed using Proteomics data from three different lifecycle stage of *T. brucei* (Dejung et al., 2016). Each transcript with non-coding potential defined in part 1) and 2) and not encoding any peptides or encoding solely non-unique peptides in part 3) were classified a Ksplice noncoding RNA genes.

#### (1) Coding potential calculator (CPC2)

The longest isoform of each Ksplice genes were used for CPC2 analysis. CPC2 (Kang et al., 2017) discriminates coding and non-coding DNA sequences based on four intrinsic features: Fickett TETSCODE socre, open reading frame (ORF) length, ORF integrity and isoelectric point (pI). The Fickett TESTCODE score was calculated from the weighted nucleotide frequency of the full-length transcript whereas the ORF length, ORF integrity and pI were calculated from the longest putative ORF identified in each gene. A CPC2 score below 0.5 defined a transcript as non-coding gene, whereas a CPC2 score ≥ 0.5 describes a transcript as protein-coding gene.

#### (2) Ribosome profiling

*T. brucei* ribosome profiling data (Vasquez et al., 2014) was re-analyzed using our merged genome annotation that consisted of the annotated protein-coding genes from TriTrypDB and our new annotated Ksplice noncoding genes (major isoforms). Quantification and statistical analysis were performed as described in Vasquez *et al*. (Vasquez et al., 2014). A *T. brucei* transcript was defined to be interacting productively with ribosomes if its translation efficacy score was ≥ 1 (translation efficacy = TE = RPKM of ribosome profiling / RPKM of RNA-seq). Inversely, A *T. brucei* transcript was defined to be not interacting with ribosomes if its translation efficacy score was ≤ 0.2857, meaning its transcript levels (RNA-seq data in RPKM) was 3.5x higher than its level of association with *T. brucei* ribosomes. And a *T. brucei* transcript with TE score in between (0.2857 < TE < 1) was defined to have low or few interaction with *T. brucei* ribosome. Additionally, as in Vazquez et al. (Vasquez et al., 2014), we investigated the 5’ end periodicity of mapped reads of both coding and putative non-coding genes. For Figure supplement 6, for each gene, the number of reads mapping to each frame of translation (represented as +0, +1 and +2) was calculated, and the frame with the highest number of mapped reads was determined. A p-value indicating the likelihood of periodicity was calculated by a binomial test on the frame with the highest number of mapped reads under the null-hypothesis that this number should be equal to ½ of all reads mapped to that gene.

#### (3) Proteomics

The mass spectrometry proteomics data from Dejung et al. (Dejung et al., 2016) was analyzed following the author’s methodology with some modifications using MaxQuant version 1.6.0.1 (Cox and Mann, 2008) and searching against our Ksplice protein database. Our Ksplice protein database is composed by 3 set of proteins – protein-coding genes from TryTrypDB (version 33, 10019 entries, excluding protein-coding genes with internal codon stop), putative proteins originated from the Ksplice new gene sequences (2003 Ksplice new genes + 72 Ksplice Kolev ncRNAs) and putative proteins originated from intergenic region sequences of *T. brucei* genome. Intergenic region sequences were selected to have on average the same size and number of sequences than Ksplice new genes. All Putative protein sequences (enclosed by a start and stop codon, with a minimum of 7 amino acids and a maximum of 4600 Da) originating from Ksplice new genes or intergenic regions of *T. brucei* genome were extracted in order of the DNA sequence and from the 3 possible translation frames (excluding sequences without start codon or/and containing ambiguous base). 14261 proteins were extracted from Ksplice new genes and 28750 proteins from the selected intergenic region of *T. brucei* genome.

### Full length sequencing

To investigate the presence of full length transcripts in our RNA-seq dataset, read pairs containing the SL sequence on the forward read and a poly-A tail on the reverse read were extracted and mapped to the *T. brucei* genome as described above (in identification of Ksplice putative new noncoding genes section). For all concordant alignments (both paired reads aligned), the boundaries of the transcripts were determined by the mapping positions of the two reads. Paired-end reads are providing the accurate boundaries of *T. brucei* transcripts as reads are sequenced from the same RNA molecule.

### Differential expression of Ksplices new genes between BSF and PCF

Differential expression analysis for Ksplice new genes between bloodstream and procyclic froms was performed using our merged annotation of *T. brucei* genome (major isoform of Ksplice new genes + Tritryp protein-coding genes) and the DEseq2 package. To that end, we used our previously published transcriptomic data (Rijo-Ferreira et al., 2017) containing 13 RNA-seq samples replicate for both bloodstream and procyclic forms.

### RIT-seq analysis of Ksplice new genes

The RIT-seq data from Alsford *et al*. (Alsford et al., 2011) was re-analyzed by aligning the sequence reads against our merged annotation of *T. brucei* genome (major isoform of Ksplice new genes + Tritryp protein-coding genes). Quantification and statistical analysis were performed as described in Alsford *et al*. (Alsford et al., 2011).

### Statistical analysis

For all graphs in Figure 2C and Figure 3C-H: the results are shown as mean (SEM, n=3) and all statistical analyses are done with two-factor mixed ANOVA (two-sided). For graph in Figure 4A: the results are shown as mean (SEM, n=4) and statistical analyses are done with two-way ANOVA (two-sided). For graphs in Figure 4B and 4C: the results are shown as mean (SEM, n=4) and statistical analyses are done with Log-rank (Mantel-Cox) test.

## Acknowledgment

The authors would like to thank Marta Machado for her precious help with the *in vivo* mouse experiment of *T. brucei* infection. We also thank Helena Manso, Ana Rita Grosso and Nuno Barbosa Morais for their valuable help in computational analysis, and Marcia Triunfol for her assistance in preparing the manuscript.

## Authors’ contributions

FG, CN and LMF designed the study. FG, FB, DN, MS designed and performed the experiments. FG, DN and LMF analyzed the data and wrote the initial draft of the manuscript. FG and LMF supervised the project. All authors edited and approved the final manuscript.

## Additional Files

Table supplement 1 – 5

Figure supplement 1 – 9

References (22 - 33)

