## Supplementary Figures for "A long non-coding RNA controls parasite differentiation in African trypanosomes"

### Figure supplements

**Figure supplement 1** – Paired-end sequencing confirms gene boundaries of 250 Ksplice novel lncRNA genes.

Examples of the genome localization of Ksplice novel genes for which paired-end sequencing has confirmed their exact gene boundaries. Ksplice coding genes that overlap Tritryp protein-coding genes (https://tritrypdb.org/tritrypdb/ ; Aslett et al. 2009) (annotated in blue) are annotated in red and Ksplice novel genes are annotated in yellow. Transcripts of Ksplice novel genes identified by paired-end sequencing are shown in green.

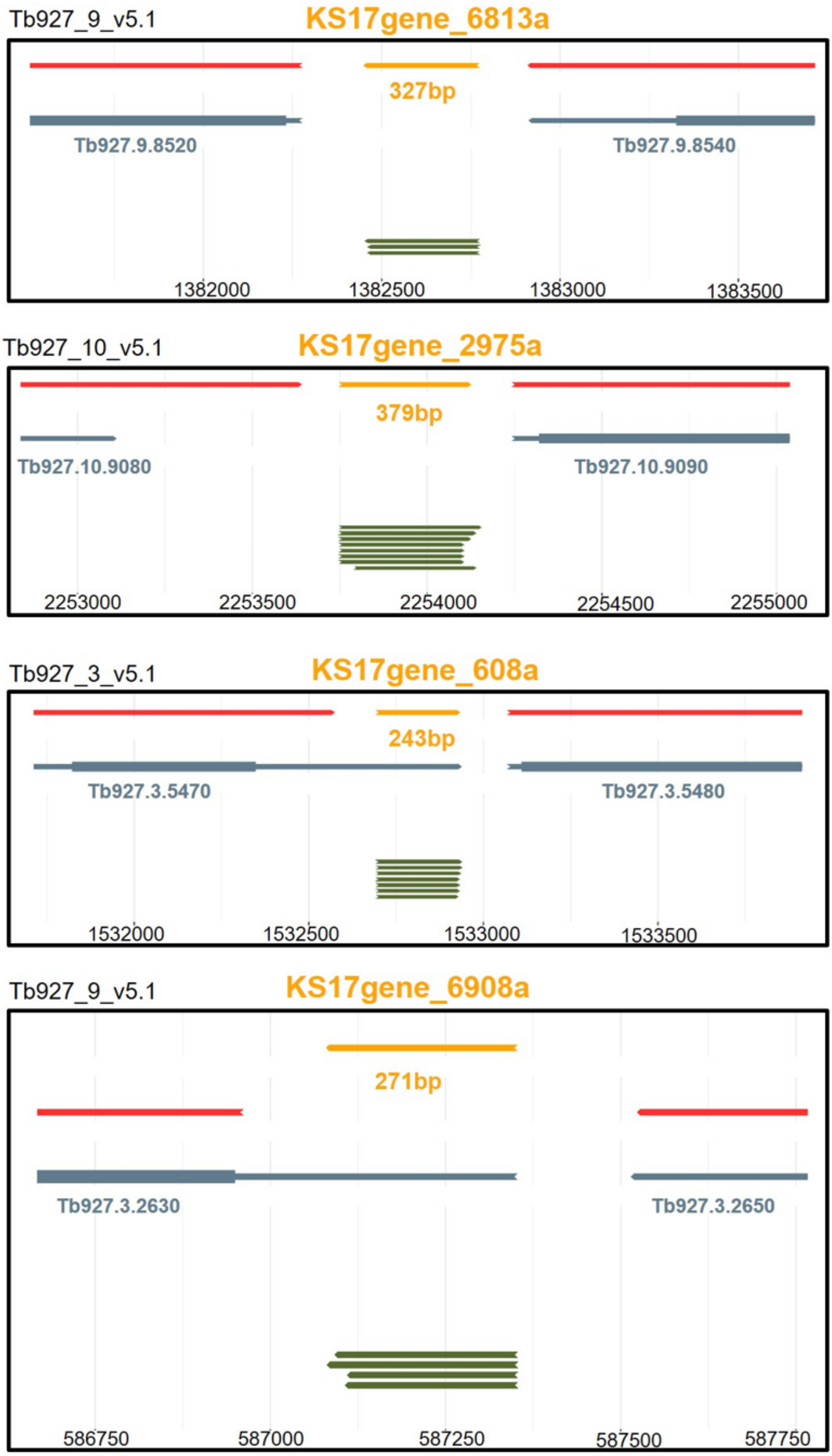

Tb927_9_v5.1
KS17gene_6813a
327bp
Tb927.9.8520
Tb927.9.8540
1382000
1382500
1383000
1383500
Tb927_10_v5.1
KS17gene_2975a
379bp
Tb927.10.9080
Tb927.10.9090
2253000
2253500
2254000
2254500
2255000
Tb927_3_v5.1
KS17gene_608a
243bp
Tb927.3.5470
Tb927.3.5480
1532000
1532500
1533000
1533500
Tb927_9_v5.1
KS17gene_6908a
271bp
Tb927.3.2630
Tb927.3.2650
586750
587000
587250
587500
587750

**Figure supplement 2** – The coding potential calculator (CPC2) score for Ksplice genes.

(A) Density plot for the coding potential calculator (CPC2) (Kang et al. 2017) score of Ksplice overlapping annotated protein-coding genes (blue curve) and Ksplice putative new genes (orange curve). Coding potential calculator score <0.5 is considered as non-coding DNA sequence. (B) Piechart showing the percentage of coding (grey) and non-coding (orange) Ksplice new genes (including 72 Kolev ncRNAs).

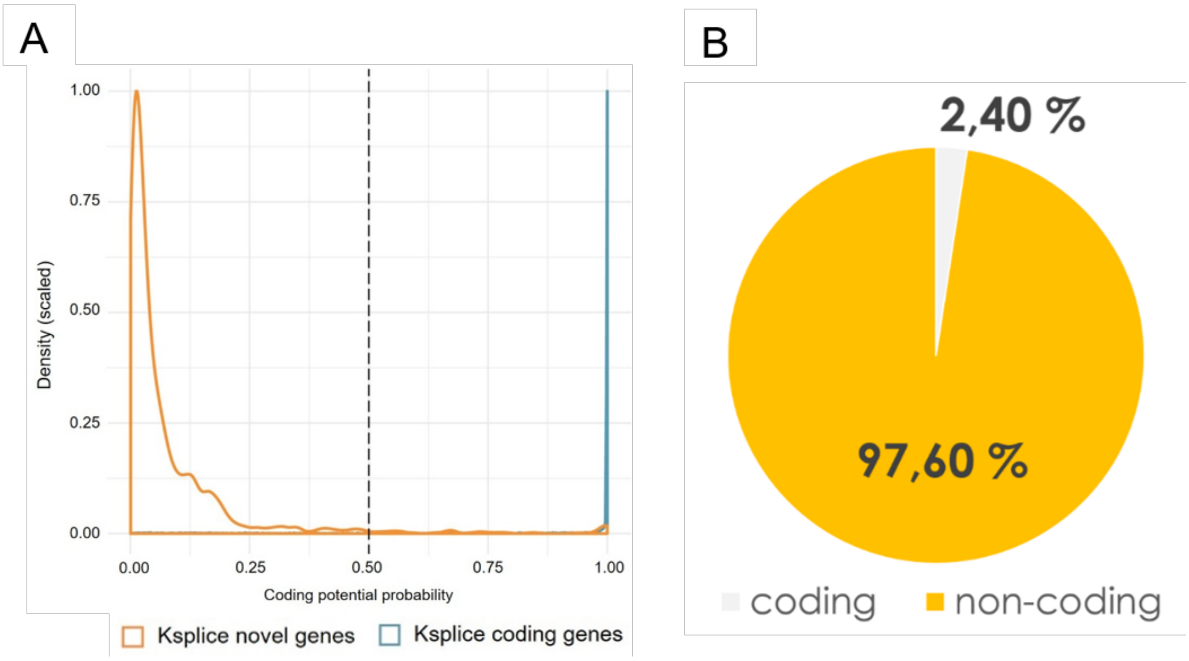

**Figure supplement 3** – Ribosome profiling reads vs RNA-seq reads for Ksplice genes.

We re-analyzed the ribosome profiling data (Vasquez et al. 2014) using our new Ksplice gene annotation of *T.brucei* genome. (A) Scatter plot showing the correlation analysis between the ribosome profiling (RF) and RNA-seq (RNA) reads of Ksplice overlapping Trytrip protein-coding genes (blue) and Ksplice new genes (orange). Pearsons correlation ($R^2$) is indicated on each plot and for each Ksplice gene types. (B) Density plot of the ribosome profiling (plain line) and RNA-seq reads (dotted line), in log10. Kpslice coding genes are shown in blue and Ksplice novel genes in orange.

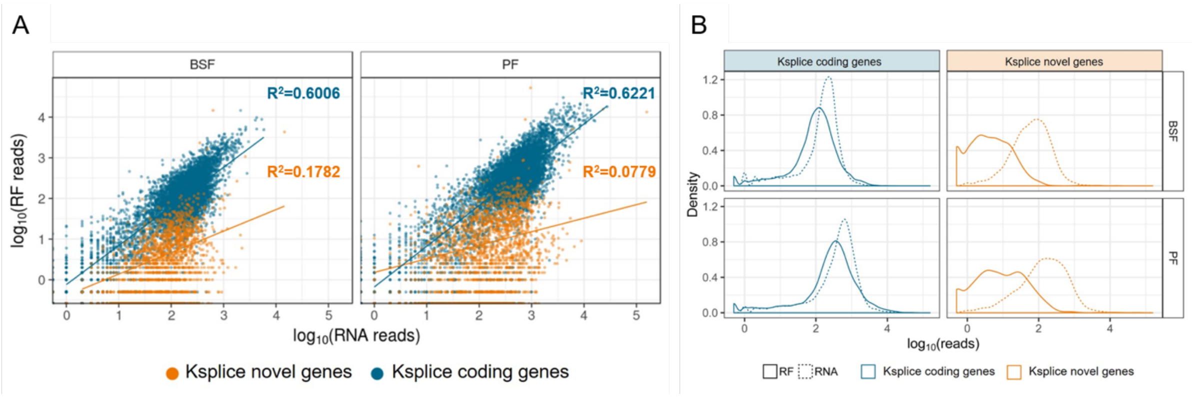

**Figure supplement 4** – Translation efficiency (TE) of Ksplice overlapping annotated genes and Ksplice novel genes.

(A) Cumulative curve of the translation efficiency (TE) for Ksplice overlapping annotated protein coding genes (blue line) and Kpslice putative new genes (orange line). Solid lines are for TE calculated in bloodstream forms and dashed lines are for TE calculated in Procyclic forms. Translation efficiency (TE) is calculated as the ratio between the reads per kilobase million (RPKM) of each transcript associated to ribosomes (Ribosome profiling) and the RPKM of that transcript in the transcriptome (RNA-seq). Transcripts with no RNA reads in RNAseq are not included in the analysis because TE=RF/RNAseq cannot be divided by 0. (B) Distribution of p-values for the frame enrichment test. No significant differences between Ksplice genes either or not overlapping with a CDS are observed for RNA levels, for ribosome footprints (RF); genes overlapping with a CDS tend to have lower p-values. (C) Numbers represent the frame containing the highest number of reads. A star (*) indicates significant enrichment.

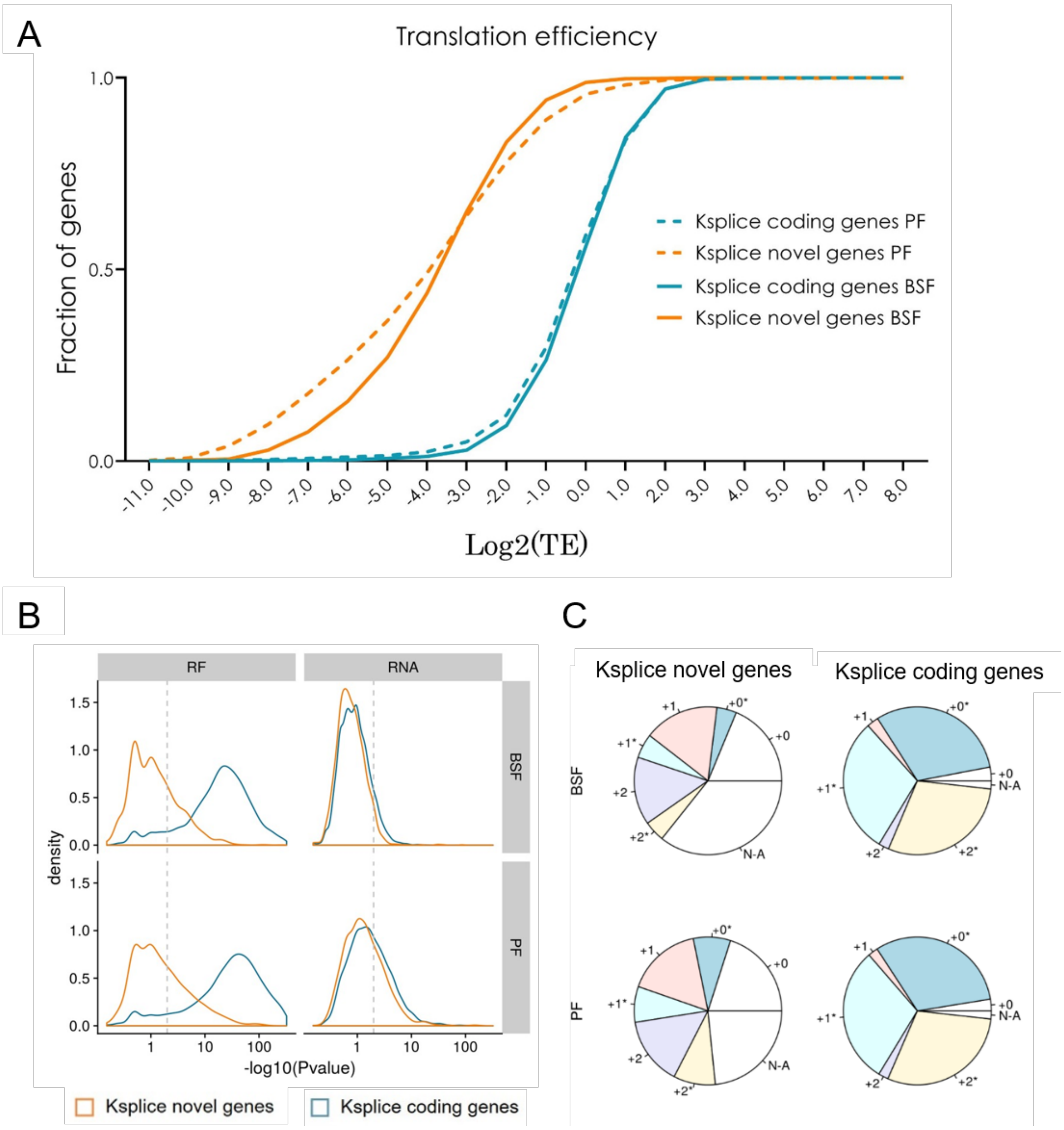
A
Translation efficiency
Fraction of genes
1.0
0.5
0.0
-11.0 -10.0 -9.0 -8.0 -7.0 -6.0 -5.0 -4.0 -3.0 -2.0 -1.0 0.0 1.0 2.0 3.0 4.0 5.0 6.0 7.0 8.0
Log2(TE)
Ksplice coding genes PF
Ksplice novel genes PF
Ksplice coding genes BSF
Ksplice novel genes BSF
B
RF
RNA
BSF
PF
density
1.5
1.0
0.5
0.0
1 10 100
-log10(Pvalue)
Ksplice novel genes
Ksplice coding genes
C
Ksplice novel genes
Ksplice coding genes
BSF
PF
+0*
+0
+1
+1*
+2
+2*
N-A

**Figure supplement 5** – Proteomic analysis for Ksplice genes.

(A) Proteomics data from dejung *et al.* (Dejung et al. 2016) was used to search for proteins/peptides translated from Trytrip annotated protein-coding genes (blue), Ksplice putative new genes (orange) and non-transcribed intergenic regions of *T. brucei* genome (grey). Putative peptides were generated *in silico* in all three reading frames of Trytrip-annotated protein-coding genes (128752 peptides), Ksplice putative new genes (14261 peptides) and non-transcribed intergenic regions of *T. brucei* genome (28750 peptides). The identification of potential proteins in each gene category was considered if at least 2 peptides were revealed by the Proteomics data. (B) The same as panel A except that putative peptides were generated from the main frame of Trytrip-annotated protein-coding genes (10019 peptides).

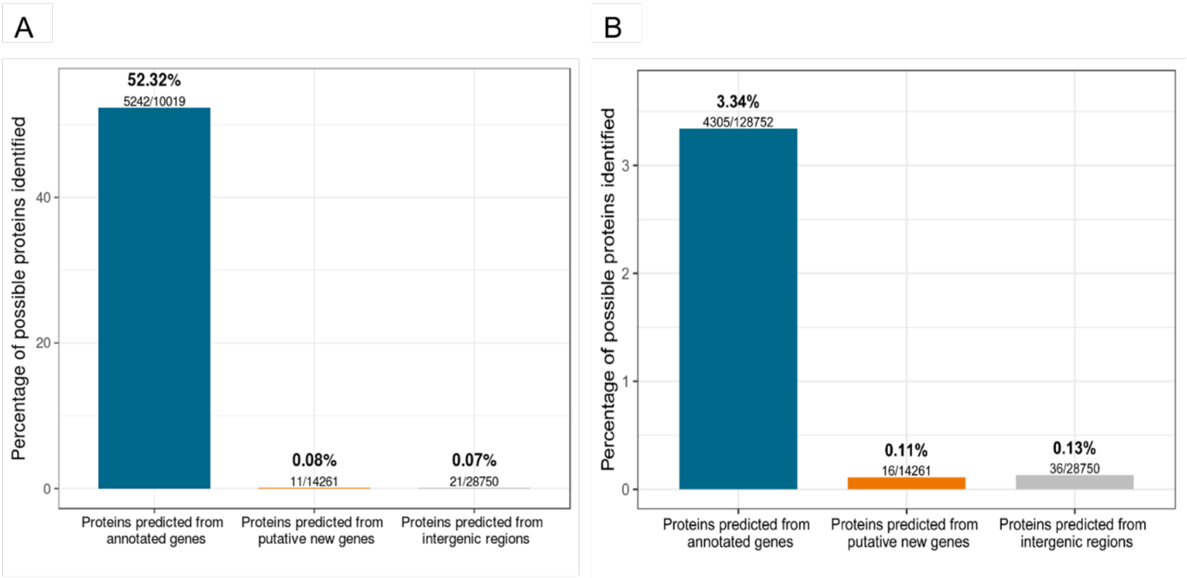

**Figure supplement 6**: – Genome location of Ksplice lncRNA genes.

(A) Genomic location and percentage of Ksplice lncRNA genes overlapping 5’UTR, 3’UTR, 5’UTR/3’UTR, or intergenic regions of *T. brucei* genome and Ksplice lncRNA genes antisense of TriTryp annotated genes.

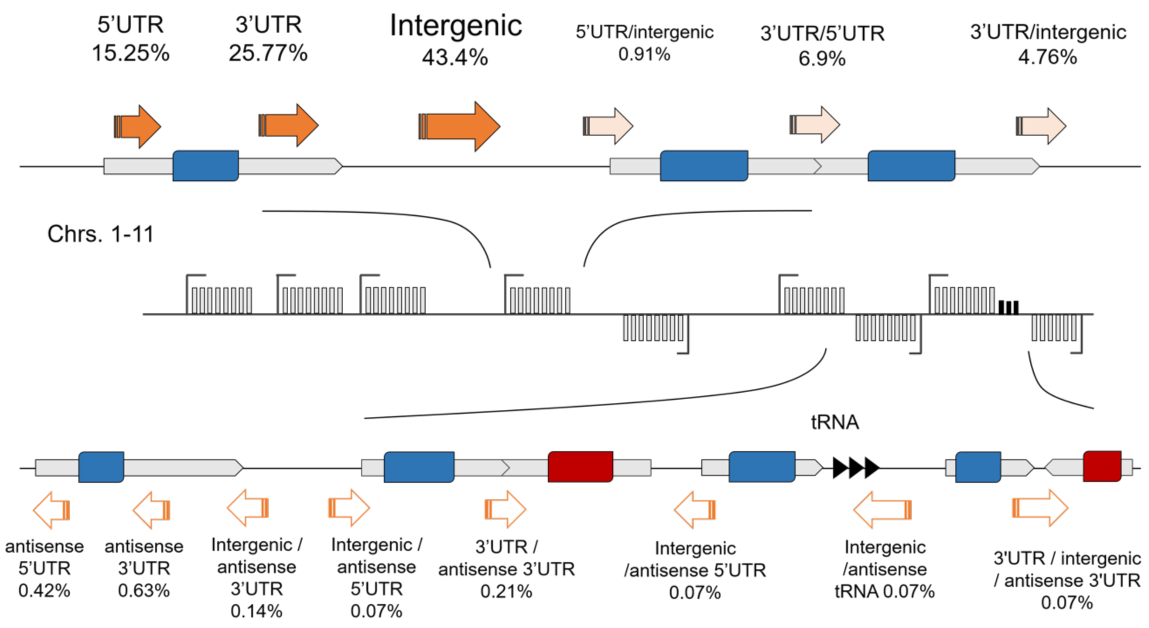

**Figure supplement 7** – Length, GC content and expression of Ksplice putative new genes compared to Ksplice overlapping annotated genes.

General characteristics of putative noncoding RNAs. (A) Distribution of the lengths of putative Ksplice putative new genes (orange) and overlapping annotated coding genes (blue). (B) GC content of putative Ksplice noncoding RNAs and overlapping annotated coding genes. (C) Distribution of mean expressions of putative Ksplice noncoding RNAs and overlapping annotated coding genes.

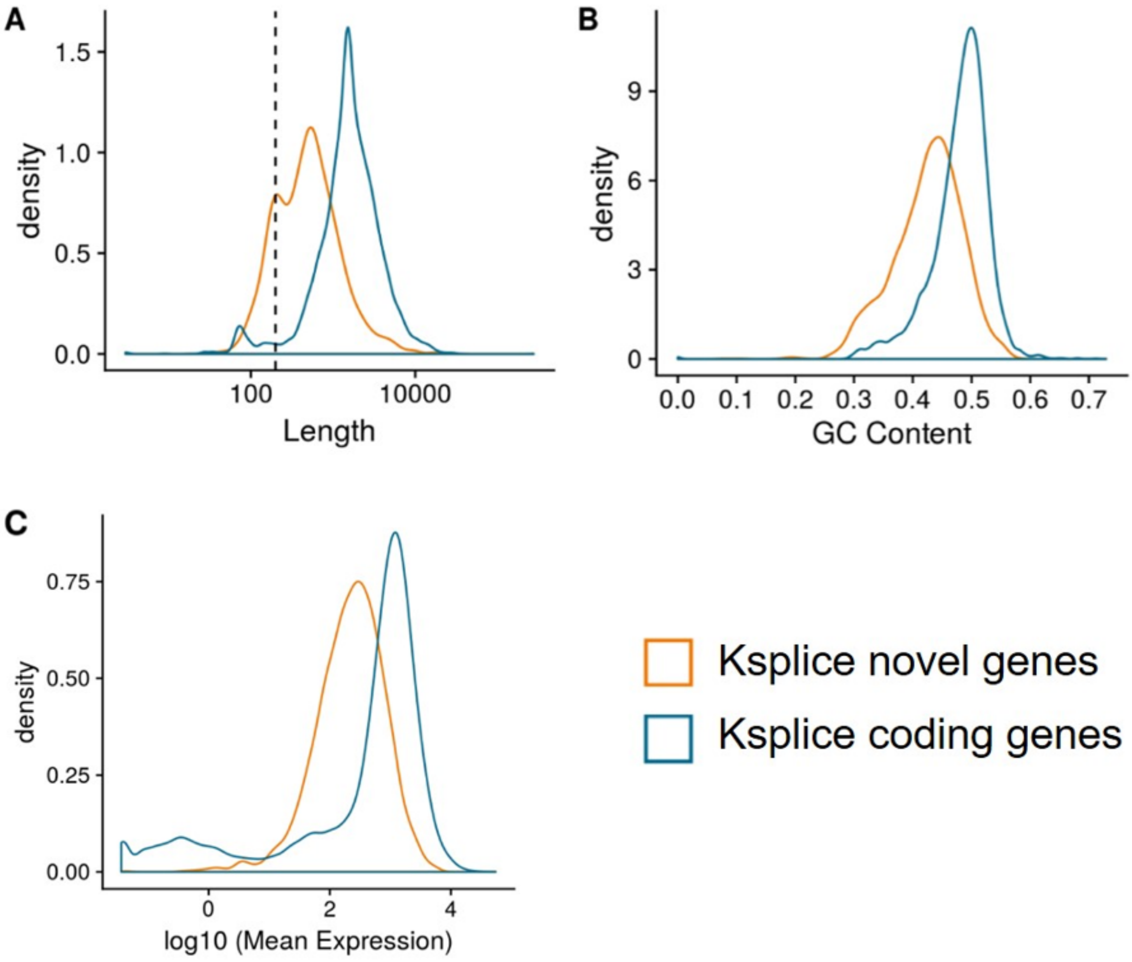

**Figure supplement 8** – Splice acceptor and polyadenylation site motifs of Ksplice genes

Splice acceptor site and polyadenylation site motifs for transcripts identified by Ksplice that either overlap with previously annotated protein-coding genes or encode for novel genes.

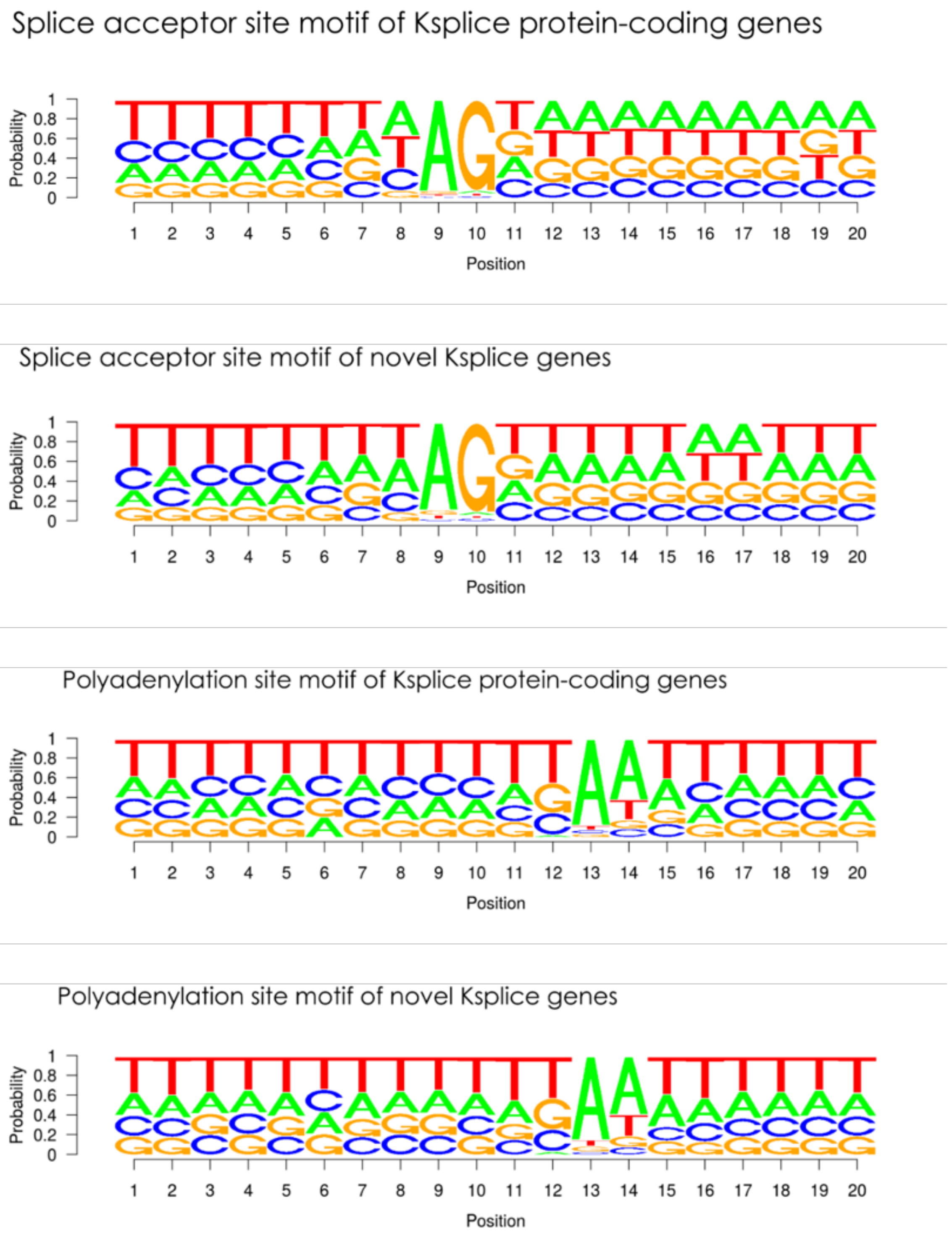

**Figure supplement 9** – Differential expression of Ksplice genes between BSF and PCF.

Differential expression of Ksplice genes between BSF and PCF our previously published transcriptomic data (Rijo-Ferreira et al. 2017) (A) Distribution of mean expression levels of Ksplice overlapping protein-coding genes (blue) and Ksplice putative new genes (orange). (B) Volcano plot of differential expression for Ksplice overlapping protein-coding genes (blue) and Ksplice putative new genes (orange). (C) MA-plot of differential expression for Ksplice overlapping protein-coding genes (blue) and Ksplice putative new genes (orange). (D) Density plot of differentially expressed genes: Ksplice overlapping protein-coding genes (blue) and Ksplice putative new genes (orange).

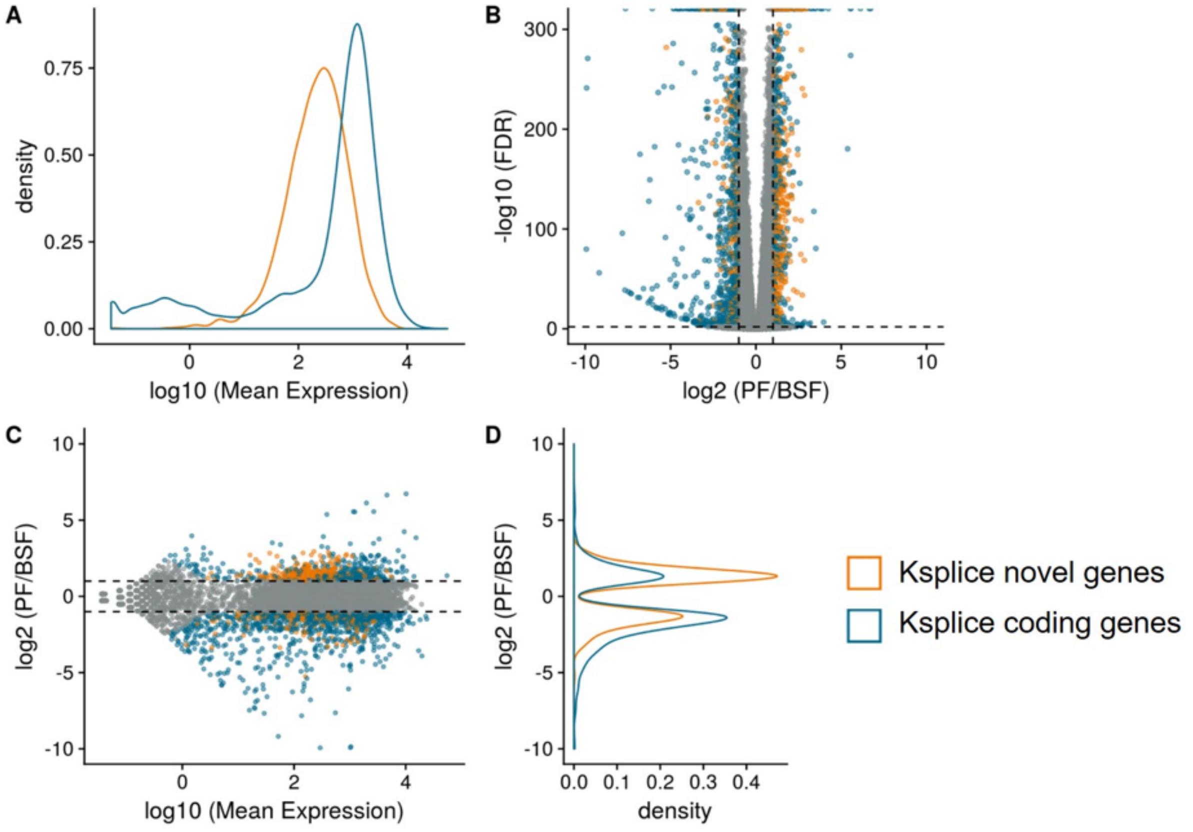
